# A Practical Approach for Optimizing Off-axis Telecentric Digital Holographic Microscope Design

**DOI:** 10.1101/2022.07.19.500723

**Authors:** Lei Jin, Ziyang Yu, Aaron Au, Christopher M. Yip

## Abstract

Digital holographic microscopy (DHM) has become an attractive imaging tool for the analysis of living cells and histological tissues. The telecentric DHM (TDHM) is a configuration of DHM that lightens the computation load with *a priori* aberration corrections. However, TDHM requires a well-aligned optical pipeline to optimize its resolution and image quality (IQ), which has traditionally complicated the alignment process. Further deriving from the optical interference functions, we offer a set of methodologies to simplify TDHM design and alignment by determining the optimal +1 order position, which depends on the object-reference beam angle and the interference plane rotation angle. The methods are then experimentally tested and verified on a TDHM system by imaging living HeLa cells in suspension.

## 1. Introduction

Off-axis digital holographic microscopy (off-axis DHM) has been of keen interest in applications ranging from particle tracking [1], microelectromechanical systems characterization [2], to live-cell imaging [3] due to its ability to acquire quantitative phase information and infer the thickness [4] and the refraction index [5] of samples from a single exposure. To recover robust object phase information from the hologram, a conventional off-axis DHM system must overcome parabolic phase aberrations introduced by the microscope objective (MO) lens [6]. Several computational techniques have been proposed to correct some aberrations *a posteriori*, such as Zernike polynomial [7] and principal component analysis (PCA) [8] at the expense of additional compute time. Recent advancement of Telecentric Digital Holographic Microscopy (TDHM) promises *a priori* phase aberration compensation by utilizing a tube lens to convert MO-induced spherical wavefront back to a planar wavefront at the camera sensor [9,10]. This optical configuration bypasses the computation cost and image quality (IQ) deterioration [11] of numerical aberration removal.

As a microscopy technique with a sub-wavelength axial resolution, TDHM is highly sensitive to optical alignment differences [12]; actuation of a single optical element may incur alteration to multiple control parameters, consequently manifesting into the final IQ [13,14]. Such a challenge is evident in the optimization for Fourier +1 order position [15,16], a key factor concerning spatial bandwidth in the filtered hologram and conductively [17], IQ of the resultant image [18]. Where a common practice to fine-tune the position of +1 order is to adjust the near-sensor beam splitter (*beam splitter 2* in **Fig. 1**) in the spatial rotational axes and actuate the preceding mirrors along the optical path [19,20], this method suffers from unwanted complexity in both the optical architecture and the adjustment procedure.

**Fig. 1.**
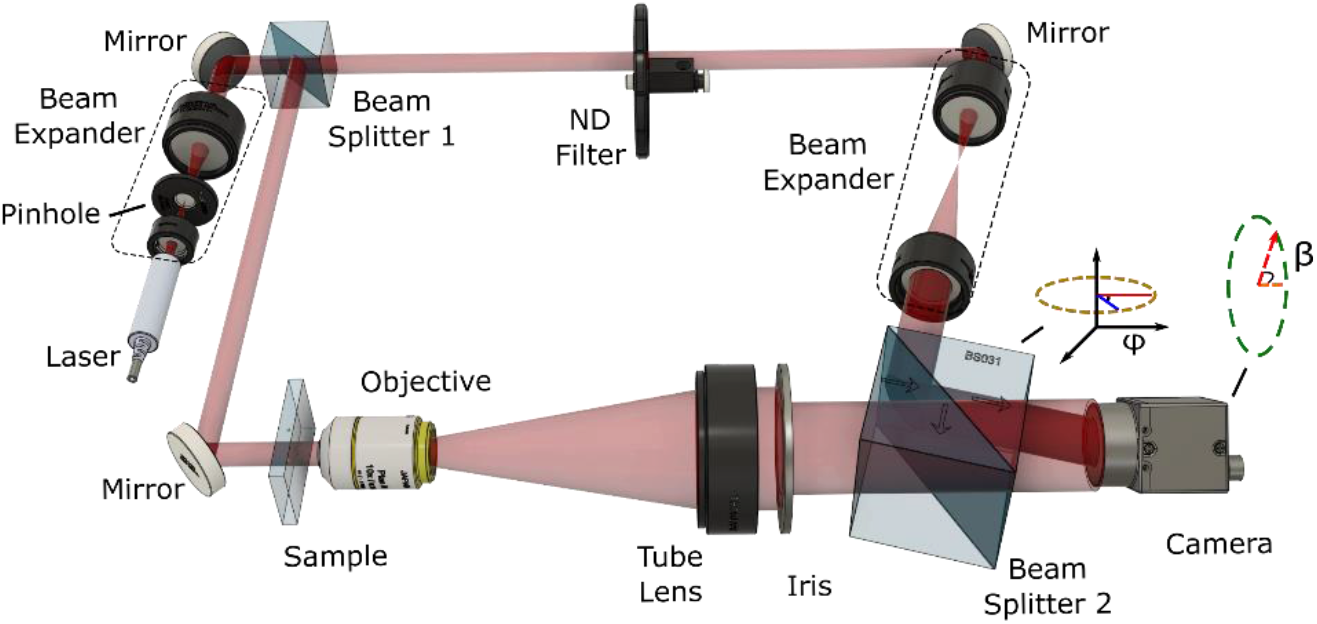
Schematic of TDHM prototype. φ represents the object-reference angle (the object and reference beams are aligned in the same horizontal plane), and β represents the rotation angle of the camera relative to the optical axis, which will be analyzed in detail in the rest of the laboratory note.

Deriving from existing mathematical models [21–24], we constructed and validated a set of comprehensive methodologies to control +1 order positioning and filtering of TDHM. This led to a design of a TDHM that isolates the control of the +1 order position to two separate one-dimensional rotations rather than the common optical train. In addition, a relationship was established between camera resolution (in pixels) and the corresponding optimal Fourier filter size. The methods presented in this laboratory note simplify the process of positioning the +1 order and improve spatial bandwidth, culminating to superior IQ.

## 2. Design Fundamental

### A. Microscope Setup

To test and validate the above approaches, a TDHM system was constructed based on the Mach-Zehnder interferometer. As illustrated in **Fig. 1**, the beam emitted by the laser (CUBE 640-40C, Coherent Inc.) is split by a beamsplitter (BS013, Thorlabs Inc.) and into the object and reference beams after passing through a beam expander (AC254-030-A-ML, AC254-060-A-ML, Thorlabs Inc.) fitted with a pinhole unit (P50K, Thorlabs Inc.). The reference beam is attenuated by an ND filter (NDC-50C-4M-A, Thorlabs Inc.) and enlarged 10x by a pair of lenses (AC254-030-A-ML, AC254-300-A-ML, Thorlabs Inc.) while the object beam passes through a MO (UPLanSApo, 20x/0.75, Olympus Inc.) and a tube lens (AC508-180-A-ML, Thorlabs Inc.). The holographic interference pattern is generated when the reference and object beams are redirected using another beamsplitter (BS031, Thorlabs Inc.) to superpose at the sensor of the CMOS camera (BFS-U3-120S4M-CS, FLIR).

On the camera sensor, the intensity projection of the interference between the object wave ***O*** and the reference wave ***R*** is described by

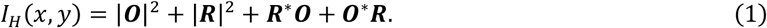

The terms in **Eq. 1** represent the zero-order (|***O***|^2^ + |***R***|^2^), +1 order (***O***^***∗***^***R***), and the conjugate -1 order (***R***^***∗***^***O***), which can be visualized in **Fig. 3 (d)**. The +1 order is filtered by either a circular or rectangular filter mask to obtain the complex optic field of the imaging samples. The filter size and position directly concern the resolution and IQ of the result image [25], as the next two sub-sections attempt to illustrate. The complex field is then reconstructed into the image plane to correct anamorphism by the angular spectrum method [26]

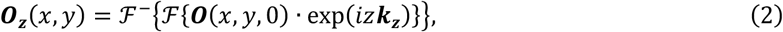

where ***O***_***z***_(*x, y*) is the complex optic field on the original plane, z represents the reconstruction distance and ***k***_***z***_ represents propagation wave vector in axial (*z*) direction. Compared to Rayleigh Sommerfeld [27] and Fresnel [28] diffraction theory-derived reconstruction methods, the angular spectrum method offers a shorter processing time with better accuracy [29]. Finally, the result phase map is recovered from the reconstructed field as

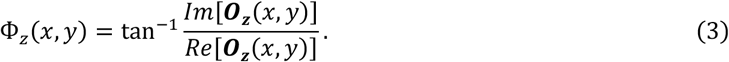

**Fig. 2.**
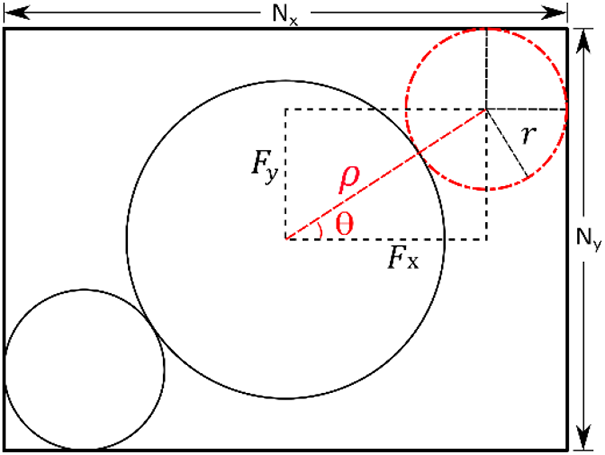
Fourier spectrum of the hologram. The shape and ideal size of diffraction orders are illustrated. For TDHM, the +1 order is near-circular, depending on wavefront curvature. The corresponding FFT filter size is described by *r*, and its position is represented in polar coordinates (*ρ, θ*) and Cartesian coordinates (*F*_*x*_, *F*_*y*_). The size of zero-order is double of +1 order [25].

**Fig. 3.**
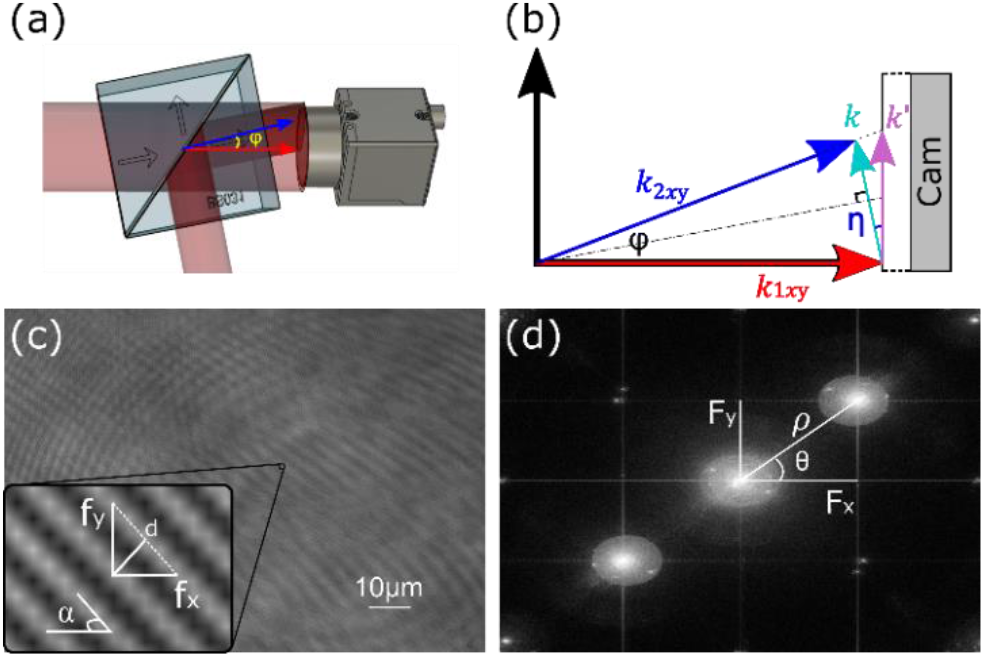
Interference from the difference in beam angles. Two beams meet at the cube and then superpose at the camera sensor, forming beam angle *φ*. The phase component can be described by wave vector ***k***. The projection of the resulting interference wave vector ***k****’* should be very close to ***k*** since the residual angle *η* is close to 0° due to the small *φ* (<10°). The hologram (c) is obtained directly from the digital camera, and (d) is the resulting Fourier spectrum, where *f*_*x*_, *f*_*y*,_ and d represent the interference fringe sizes, and (*F*_*x*_, *F*_*y*_), (*ρ, θ*) represent the corresponding +1 order position in the Fourier spectrum in Cartesian and polar coordinates, respectively.

The phrase Φ_*z*_(*x, y*) represents the optic path length difference between the object beam and the reference beam, which equals the multiplication between the physical propagation distance and the comprehensive refractive index. Combining the above methods, the processing pipeline has been extensively implemented on many DHM systems [15] and is also employed by our TDHM setup, where the processing steps are executed in series to obtain a phase map.

### B. Fourier Filter Optimization

The selection and application of an appropriate FFT filter is a critical step in the TDHM processing pipeline. With the zero-order intensity filtered out, the FFT filter mask should maintain good coverage over the high-frequency components of the hologram. Due to TDHM’s use of a tube lens for spherical-to-planar wavefront conversion, the shape of the +1 order would approach a perfect circle (**Fig. 2**), making it appropriate for the use of a circular filter [25]. Maximizing the filter size allows high spatial bandwidth and the subsequent high lateral resolution in the resulting image [21].

Expressed below are a set of equations pertinent to the Fourier filter of **Fig. 2** (pixel unit in Fourier space),

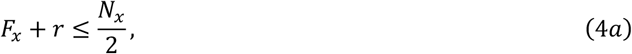

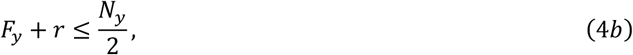

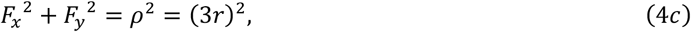

where (*F*_*x*_, *F*_*y*_) and (*ρ, θ*) represent the position of +1 order in Cartesian and polar coordinates, respectively. The *r* represents the radius of the circle FFT filter; *N*_*x*_ and *N*_*y*_ represent the number of pixels of the camera sensor in the *x* and *y* directions. The size of zero-order is double of +1 order, according to [25]. Substituting **Eq. 4a** and **Eq. 4b** into **Eq. 4c**, the radius of the filter can be described by (pixel unit in Fourier space)

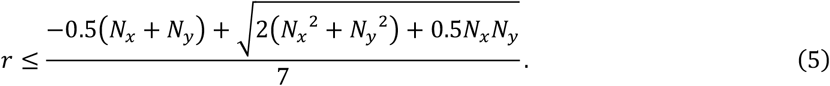

where the maximized filter radius only depends on the pixel numbers of the camera sensor. **Eq. 5** expands upon the work of Sánchez-Ortiga *et al*. [25] to include the application on cameras with non-square sensors, which are more common. The optimized +1 order position is obtained in polar coordinates as (pixel unit in Fourier space),

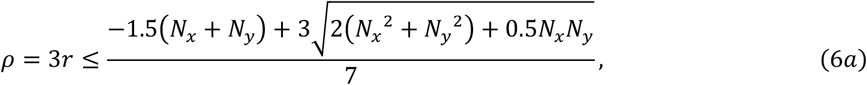

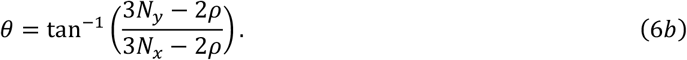

In Cartesian coordinates, the optimized +1 order position is described as (pixel unit in Fourier space),

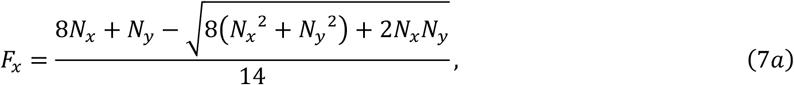

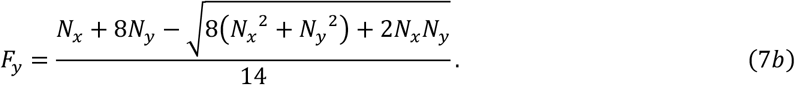

It can be reasonably observed from **Eq. 7** that the +1 order position with maximum bandwidth is unique to each combination of camera pixel dimensions. It is worth mentioning that the optimized +1 order is not always on the image diagonal as it is the case for square sensors; **Eq. 6** and **Eq. 7** can otherwise provide the optimized +1 order position for TDHM systems employing non-square sensors. For instance, our camera (BFS-U3-120S4M- CS, FLIR) has a non-square camera resolution of 4000 × 3000, and its optimized +1 order position is (1707, 33.04°) (Polar coordinates) and (1431, 931) (Cartesian coordinates) in pixel units in Fourier space, which is off the sensor diagonal.

### C. Order Position Alignment

This section sets out to detail a simpler method to control the position of the +1 order. Instead of aligning the object and reference beams using a 6-degree of freedom optical system, we propose a simplified alignment method that utilizes two independent, one-degree rotational parameters (the beam angle and the camera rotation angle) based on Goodman’s derivation of optical interference [21].

The alignment process begins by placing the object and reference beam onto the same plane to eliminate the effect introduced by fringe pattern tilt angle α. As the object and reference beams intersect on the sensor, **Fig. 3a** and **Fig. 3b**, the consequent spatial beam angle, φ, dictates two attributes to the interference fringe pattern: the fringe tilt angle, α, as well as the interference fringe size, *d* (**Fig. 3c**). To characterize the relationship between beam angle and the position of the +1 order, we can first determine how interference patterns on a camera sensor affect the position of +1 order in Fourier space. With a simple geometric relationship [21,22], as shown in **Fig. 3**, the distance between the interference fringes is described by:

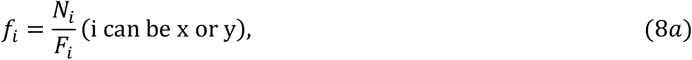

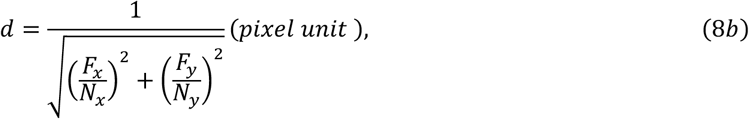

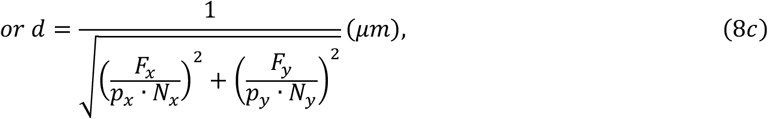

where *f*_*x*_, *f*_*y*_ represent the interference fringe distances, and N_i_ is the pixel number of the camera in x and y direction, (*p*_*x*_, *p*_*y*_) are the pixel size of the camera in μm. According to the geometric layout (**Fig. 3 (a)** and **(b)**), the fringe size (*d*) is described by the camera pixel numbers and position of the first order in the spectrum.

Assuming the optic wave of the DHM is propagating in a dielectric medium whose magnetic and electric permittivity are constant throughout the region of propagation and independent of the direction of the polarization, the effect of the medium can be neglected when we only consider the image intensity contrast on the camera sensor. The interference between the object and the reference beam can be represented by:

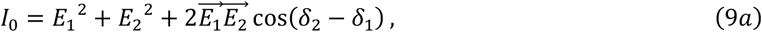

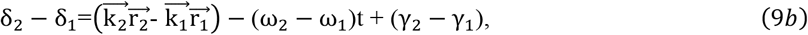

where *E*_*i*_ is the amplitude of the two beams, 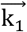 and 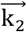 are wave-vectors, *ω* represents the wave frequency, *γ* represents their initial phase, and *δ*_*1*_ and *δ*_*2*_ represent their comprehensive phase. In this case, as the two beams are emitted by the same laser source, *(ω*_*2*_*-ω*_*1*_) and *(γ*_*2*_*- γ*_*1*_) are both equal to zero. Using ND filters and polarizers, the intensity of beams can be equalized (*E*_*1*_*=E*_*2*_*=E*_*0*_). The interference expression can be simplified as

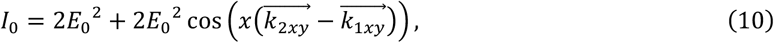

where 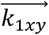 and 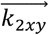 are wave vectors of the object and reference beam in the horizontal plane, respectively. Here, **Eq. 10** represents the intensity distribution of the optic field encompassed by the two beams. The interference fringes are encoded in the phase difference between the two beams 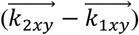. In this case, we can describe the phase difference by 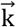 • *sin*(*φ*/2). From **Fig. 3 (b)**, we know an error is present between *k* vector and the actual *k′* vector due to the residual angle *η* (*φ/2* in this case). This error was ignored when the camera was close enough to the front cube since the beam angle (*φ*) is generally small (<10°). With this assumption, the interference pattern can be simplified as

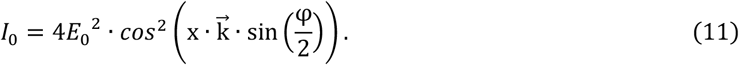

The fringe distance *d* is exactly the period of this projected intensity distribution (**Eq. 11**). The wave vector ***k*** can be described as *2π/λ*, where *λ* represents the illumination wavelength. Thus, the fringe size is described as [21]

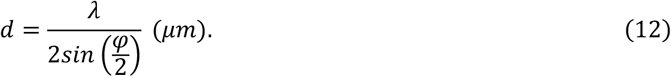

Combined with **Eq. 8b**, the beam angle can be described as

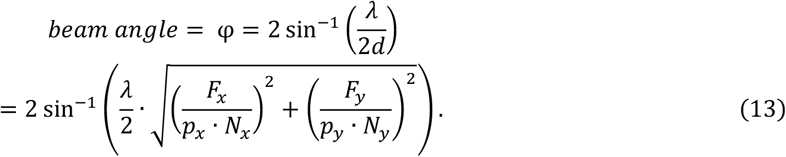

where *p*_*x*_ and *p*_*y*_ represent the pixel size (μm) of the camera in the x and y directions, respectively. **Eq. 12** illustrates that the fringe size depends on the illumination wavelength and the object-reference beam angle, which is experimentally verified in the next section; the optimized beam angle can be subsequently calculated with the knowledge of the +1 order position. Beam angle variance influence greatly on the separation between +1 order and zero order, ρ, while having a negligible impact on order angle θ. To optimize the +1 order position, the object- reference plane can be rotated around the optical axis, otherwise equated by the rotation of the camera. The rotation angle can be obtained with **Eq. 8**, which yields

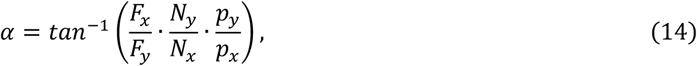

To synopsize this section, an experimental setup and process workflow were introduced, which paved the way for the calculation procedure of the optimized +1 order position. Lastly, a practical method of controlling the +1 order position by beam angle adjustment and plane (or camera) rotation was adduced. By doing so, we are able to isolate the alignment parameters to individual optical components.

## 3. Experimental Results

In this section, three experiments were conducted to verify the control methods detailed in the last section. The precision of +1 order control by either the beam angle or the rotation of the camera (or plane) was examined, and its error range was specified. In addition, the effect of various Fourier filter sizes and camera pixel sizes on TDHM system IQ was investigated.

### A. Beam angle vs. +1 order distance

A controlled experiment was set up to validate **Eq. 12**, with object-reference beam angle as the independent variable altered in each experiment trial. To satisfy the requirements, we set fringe pattern tilt angle α to 0 by placing the object beam, reference beam as well as camera on the horizontal plane as discussed in **Section 2C**. The camera was moved to two positions along the optical axis for each trial (**Fig. 4(a))** for measurement of the beam angle; it should be noted that the fringes pattern size is immutable to the axial position variation of the camera (**Fig. 5**), coinciding with the expression in **Eq. 12**. A set of datapoint containing beam angle – fringe size pairs can then be obtained across the experiment to compare with **Eq. 12** curve data.

**Fig. 4.**
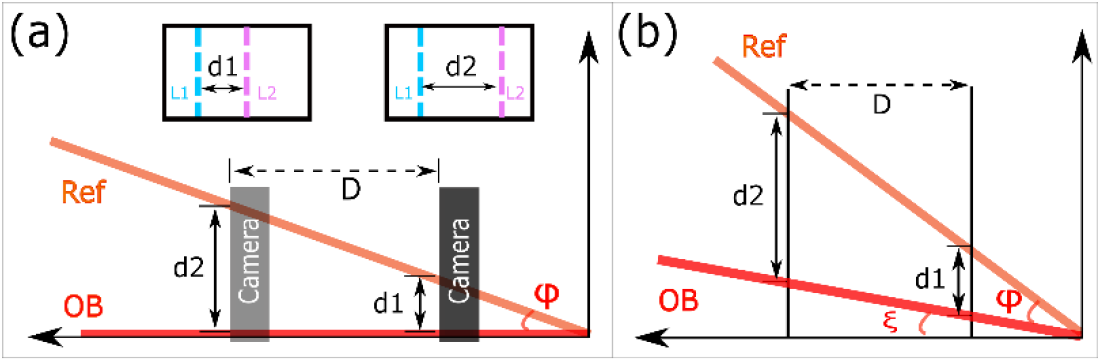
Experiment setting for beam angle. (a) Experimental setup, (b) error analysis. Setting a line pattern into the object (black dotted) and reference arms (purple solid), respectively. The projected line distances *d*_1_ and *d*_2_ can be obtained by moving the camera at a distance *D* along the optical axis.

**Fig. 5.**
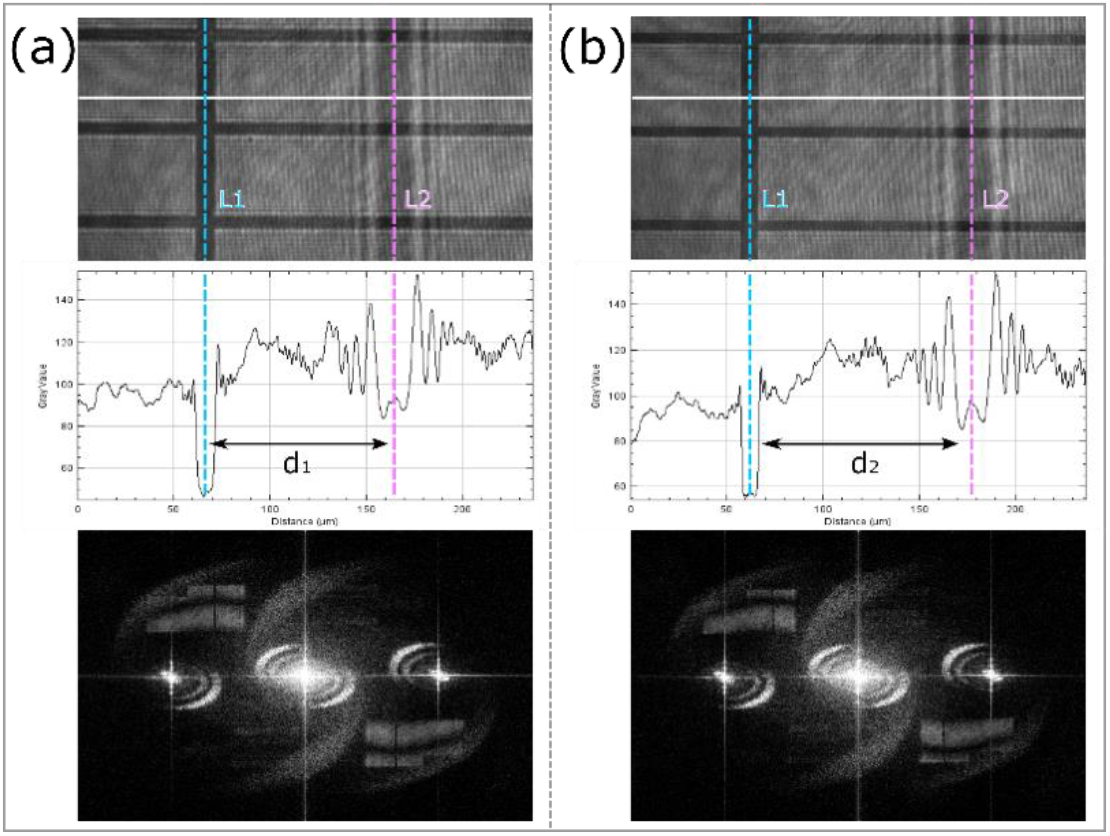
Scaling of the line pattern. The distance variance between line pattern *L*_*1*_ and *L*_*2*_ with respect to 5mm displacement of the camera (*D* = 5mm in **Fig. 4)**, before (a) and after (b). After the raw hologram is Gaussian-blurred, the line distance can be obtained from the respective intensity plot (row 2). It is noted that with the displacement of the camera, the distance between *L*_*1*_ and *L*_*2*_ has changed from *d*_*1*_ to *d*_*2*_, but their Fourier spectrums remained identical (row 3), evincing the immutability of the fringe pattern size with respect to camera displacement. This method is also used to verify the beam angle function in **Fig. 6**.

To determine the incoming beam angle, a line target orthogonal to the rotational axis was introduced into the beam paths, as illustrated in **Fig. 4(a)**. Since the beams are collimated, the projection of the line pattern is the same at any point along the path, and only the distance between the reference (Ref in **Fig. 4(a)**) and object (OB in **Fig. 4(a)**) beams changes. The beam angle (φ) can be determined by measuring the distances between the lines (*L*_*1*_, *L*_*2*_) at two different camera positions along the optical axis, as shown in **Fig. 5**, characterized by a simple geometric relationship,

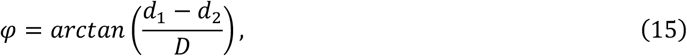

where *d*_*1*_ and *d*_*2*_ are the distances between the projected lines *L*_*1*_ and *L*_*2*_, and *D* is the displacement of the camera in **Fig. 4(a)**.

However, we must account for a small angle (ξ) between the OB path and the optical axis to compensate for misalignment, as depicted in **Fig. 4 (b**). Accordingly, when the beam angle φ is between 0° and ∼ 10°, it can be described as:

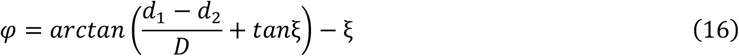

If properly aligned, the residual angle is typically less than 0.3°, and the approximation can be simplified to:

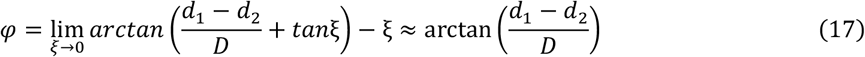

We note that **Eq. 17** can be approximated to **Eq. 15** as we assumed the residual angle (ξ) is negligible. The fringe size *d* can be obtained via **Eq. 8**(b), which requires selecting the optimized +1 position *F*_*x*_, *F*_*y*_ in the Fourier spectrum (**Fig. 5**, row 3) in addition to the knowledge of camera resolution (*N*_*x*_, *N*_*y*_) and pixel size (*p*_*x*_, *p*_*y*_). These values, coupled with their respective beam angle in each trial from **Eq. 17**, are plotted in comparison to the function curves of **Eq. 12** for verification, as illustrated in **Fig. 6**. The experimental Root-Mean-Square-Error (RMSE) for all trials is 0.131μm^2^.

**Fig. 6.**
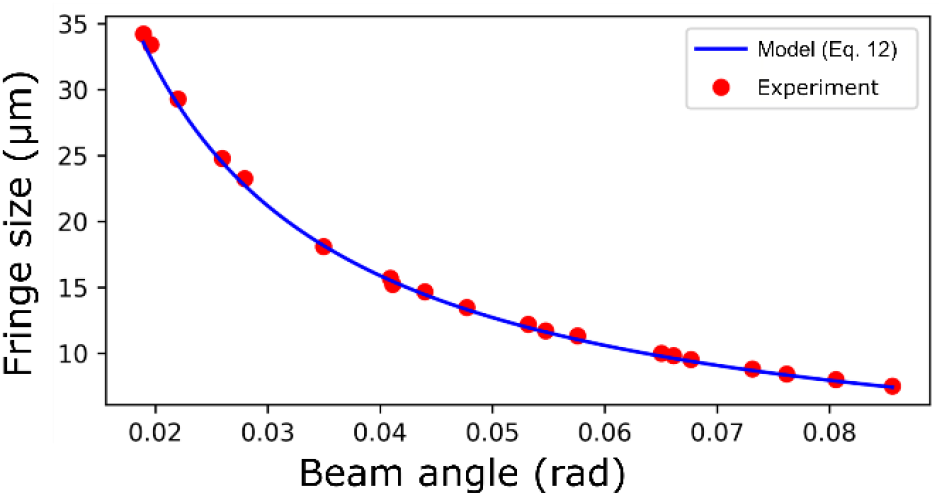
Beam angle vs. Fringe size between Eq.12 Model curve and Experiment data. The blue fitted curve is the result of **Eq. 12** with the beam angle - fringe size data points from the experiment in **Section 3A**.

### B. Camera angle vs. Fringe angle

The above experiment assumes that the object and reference beams interfere on the same plane, where the fringes pattern and corresponding first order would be orthogonal to the field of view, as in **Fig. 5**. Nevertheless, the optimal +1 order position angle *θ* should be adjusted to optimum according to **Eq. 6**, which is conventionally realized by spatially tilting the reference beam and the *beam splitter 2* until the angle of the fringes is acceptable [30]. A simpler approach would be to either rotate the object-reference plane or alternatively, the camera sensor around the optical axis to achieve the optimal order position angle *θ*, as demonstrated in **Fig. 7**.

**Fig. 7.**
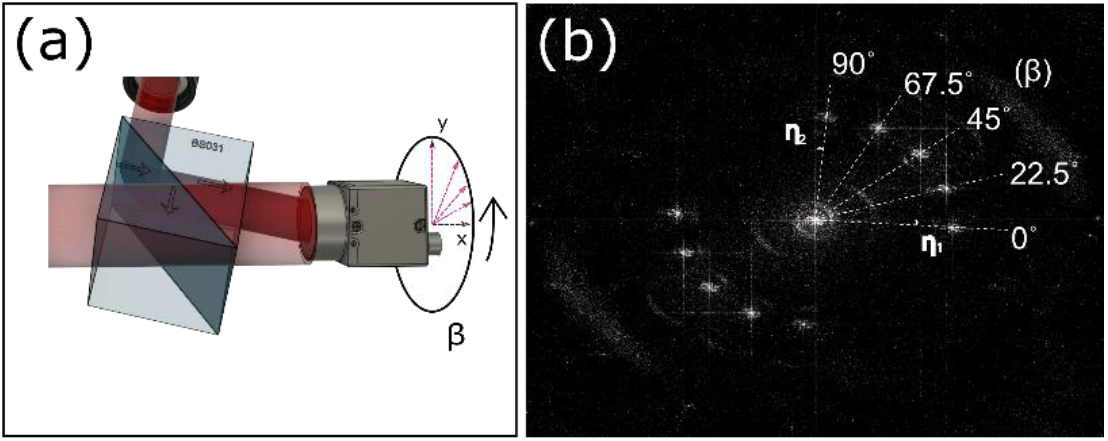
Optimization of camera rotation. (a) Schematic of the experiment where the camera is rotated around the optical axis; (b) corresponding order positions in Fourier spectrum, where β is the camera rotating angle; η_1_ and η_2_ are typical system errors mainly from the residual beam angles that are calculated as 0.2° ± 0.03°.

By rotating the camera sensor around the optical axis, the corresponding +1 order angle *θ* (0°, 22.5°, 45°, 67.5°, and 90°) can be obtained as in **Fig. 7(b**). When the camera is set at 0° or 90°, rotational errors arise in the first order of the spectrum since the object and reference beams do not perfectly interfere at the same plane, and the object beam is not perfectly parallel to the optical axis. However, the corresponding residual beam angle (0.2° ± 0.03°) can be easily obtained by **Eq. 13** and compensated with additional camera rotation. We then calculated the resulting +1 order angle, *θ*, in polar coordinates by **Eq. 6** and **Eq. 7**, which only depends on the camera camera resolution; the camera rotation angle is subsequently evaluated by **Eq. 14**.

Our CMOS camera (BFS-U3-120S4M-CS, FLIR) has a non-square sensor with camera resolution of 4000 × 3000, unlike the one used by Carl *et al*. [31], resulting in an optimal +1 order position of (1431, 931) (Cartesian coordinates) in pixel unit in Fourier space (**Eq. 7**) and the corresponding filter radius is 1707.2 (pixel unit in Fourier space). Solved with **Eq. 18**, the optimized camera rotation angle β should be 49°, and the corresponding +1 order angle θ would be 33.04°. The +1 order is located off the diagonal of the spectrum, which is also in line with the results presented in **Section 2B**.

### C. FFT filter vs. Image Quality

To verify the proposed optimization methods in the TDHM pipeline, an imaging experiment was set up to examine TDHM IQ with various beam angles and camera pixel size settings. A sample of living HeLa cells was prepared and diluted with Dulbecco’s Phosphate Buffered Saline (PBS, D1408, SIGMA Inc.). 2 ml of the mixture was injected into a petri dish (catalog#153066, Nunc Cell Culture dishes, Thermo Fisher Scientific Inc.) and left stationary for 10 min before being placed into the sample holder of TDHM without any additional labeling or treatment performed. The HeLa samples were imaged with two different beam angles settings, 2.9° and 6.6°, by an Iris 15 camera (Cam1, (Teledyne Photometrics Inc.)) and a FLIR camera (Cam2, BFS-U3-120S4M-CS, FLIR Inc), whose pixel sizes are different. We arbitrarily chose an object-reference beam angle of 2.9° on Cam 1 after yielding a usable image on the camera as **Fig. 8** (column 2); its filter size in pixel value is then mirrored onto Cam 2, which gives a corresponding beam angle of 6.6° according to **Eq. 13** and produced images as shown in **Fig. 8** (column 3). The two beam angles were then swapped and applied onto the other camera to test the extremities of the filtering condition, with the results shown in **Fig. 8** (column 1) and **Fig. 8** (column 3). The refractive index of HeLa cells was assumed to be 1.33, closing to PBS [32]. After processing the sample through the imaging pipeline described in **Section 2**, the 3D topographies of the holograms, as well as their height plots, were obtained and rendered by Fiji (Interactive 3D Surface Plot) [33], as depicted in **Fig. 8** (row 3).

**Fig. 8.**
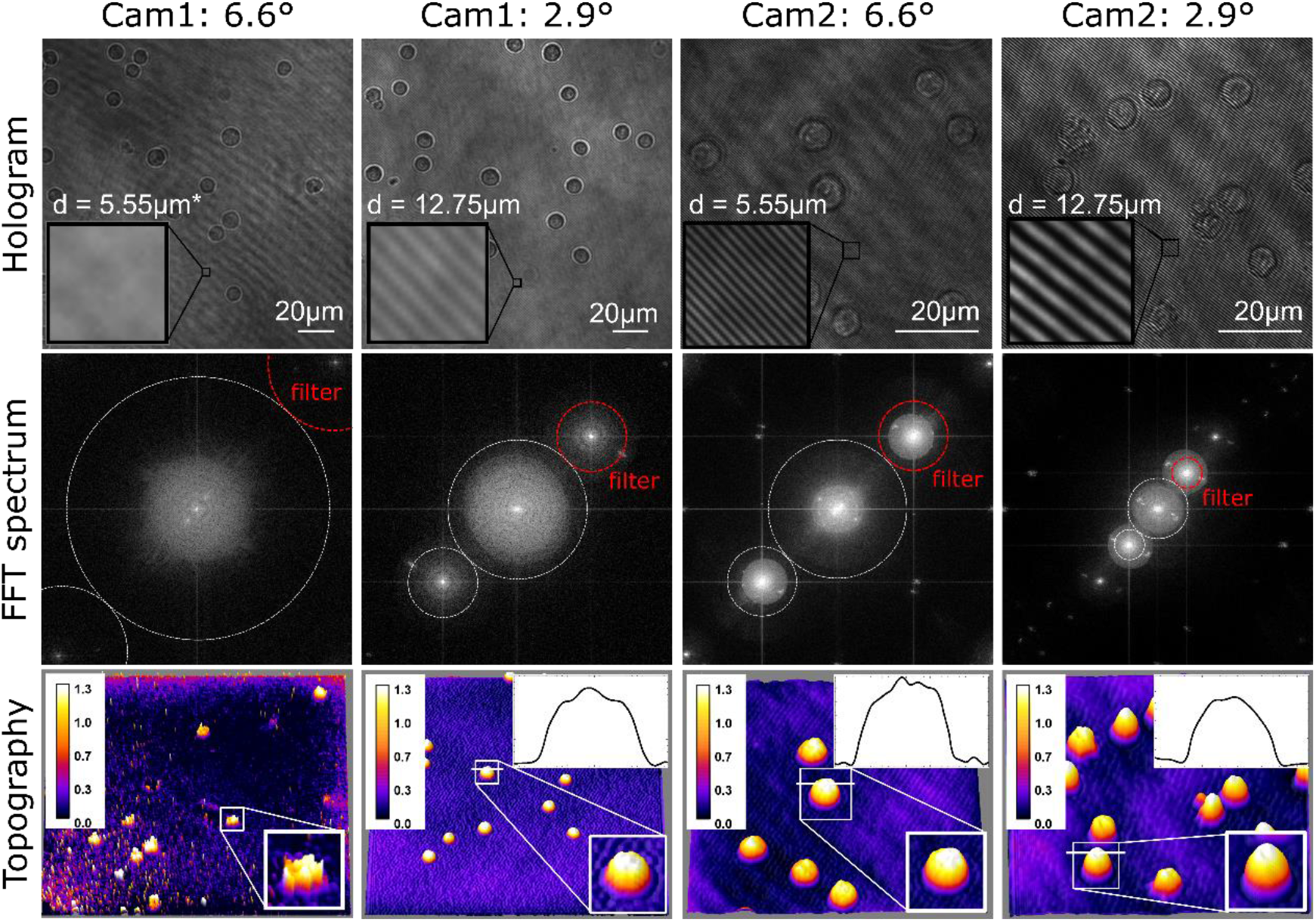
TDHM result of HeLa cells with different beam angles and camera pixel sizes. Cam 1 is Iris 15, Teledyne Photometrics Inc (pixel size: 4.25μm); Cam 2 is BFS-U3-120S4M-CS, FLIR Inc (pixel size: 1.85μm). All scale bars of various drawn lengths in row 1 are of physical length 20μm, their discrepancy due to pixel sizes difference between the two cameras. The object-reference beam angle of 2.9° and 6.6° are alternatingly applied to Cam 1 and Cam 2. The illumination wavelength is 640nm. The color scale bars in row 4 are in μm units. The refractive index of the HeLa cells was assumed to be 1.33 [32], as their 3D topographies and the height plot were obtained and plotted in Fiji, shown in row 3. The sizes of all height plots are 13μm and 1.3μm for the x and y directions, respectively. The size of the zoomed-in Region of Interest is 13 × 13 *µm*. *Fringe pattern in row 1, column 1 subfigure is not visible in the hologram since *d* = 5.55μm is less than 2 times the pixel size (8.5μm) specified in the Nyquist-Shannon sampling theorem.

According to **Eq.8(c)** and **Eq. 12**, the fringe sizes of these holograms are 5.55μm (6.6°) and 12.75μm (2.9°), respectively, for the cameras. It is noted that in one trial where the fringe size was less than the 2 × multiplier to pixel size (4.25μm) of the Iris 15 camera with 6.6° beam angle in **Fig. 8** (column 1); by Nyquist-Shannon sampling theorem, the fringe size is less than the Nyquist Rate threshold therefore not sufficient to be sampled by the pixels. This results in the virtual absence of fringe pattern details in **Fig. 8** (row1, column 1), whose +1 order was too far from the zero-order in its Fourier spectrum to produce a usable topographic image. On the other end of the spectrum, the relatively large size of the fringe pattern (12.75μm) with respect to pixel size (1.85μm) in the FLIR with a 2.9° beam angle in **Fig. 8** (column 4) only permits a small FFT filter not able to fully cover the high-frequency components without risking acquisition of zero order information, consequently losing high-frequency topography details in comparison to the other two configurations.

Several works of literature have reported on the topic of DHM operation at diffraction limit [34–36], where its resolution is mainly influenced by MO characteristics and illumination wavelength. However, the employment of an unoptimized Fourier filter inhibits its near-diffraction-limit performance. The above experiment verified the FFT filter mask needs to be sufficiently large to cover high-frequency components of the hologram for high lateral resolution without corroding into the zero order, and an optimal FFT filter (+1 order position) depends on the illumination wavelength and the camera pixel size and camera resolution.

## 4. Discussion

TDHM improves upon DHM’s aptitude in single-shot quantitative phase imaging by canceling out spherical aberrations in conventional DHM systems. However, its propensity for +1 order position sensitivity opens to high-order, multiplex effects on the IQ by adjustment of isolated optical elements, making the conventional alignment method difficult. In this laboratory note, we presented a suite of verified methodologies to mathematically locate and mask the optimized +1 order position for a given TDHM system using few parameters possible, as well as best practices to control the +1 order position and filtered bandwidth by physical alignment and processing. Its novelty arises from the fact that the alignment task can be approached with one-dimensional readily-computable adjustments to the yaw angle of *beam splitter 2* and the rotation angle on optical axis of the camera.

The proposed method is independent of the MO used in the TDHM setup due to the planar-wave illumination. Though the optical resolution and the density of TDHM are determined by the properties of MO (magnification and numerical aperture) and illumination source, the interference pattern would not be influenced. The proposed method sets out to minimize much of the resolution loss during image acquisition and phase signal processing defined by the Optical transfer function (OTF).

The method first identifies camera resolution (in pixels) as the only attribute to +1 order position (pixel units in Fourier space); a robust relation to the optimal +1 order position and circular filter geometry is consequently established, which can be implemented to various camera pixel and sensor shapes. Thirdly, the order positioning alignment of TDHM can be isolated down to two one-dimensional rotational parameters: object-reference beam angle (adjusted by *beam splitter 2* **Fig. 1**) and camera rotation angle (or object-reference plane) about the optical axis. Lastly, the expression for interference fringe pattern size can be obtained as dependent on only the beam angle and illumination source wavelength.

By using our simplified alignment approach, we verified the order positioning method by a controlled experiment to isolate and calculate beam angle in each trial; the experimental result confirms the correlation between fringe pattern size and beam angle. Equivalency between camera rotation angle and object plane rotative angle on the optical axis is confirmed in the experiment, and a more robust +1 order positioning formula is applied in polar coordinates. Lastly, we compared the IQ of images processed with different Fourier filters, demonstrating the merit of large filter size. These experimental validations come together in **Fig. 9**, where a result phase map is obtained using the computed optimal +1 order position and, consequently, a maximal Fourier filter size (1707 pixels in radius). Compared to the largest usable filters (1000 pixels in radius) in **Fig. 8** (columns 2, 3), the optimal Fourier filter encompasses more high-frequency components, preserving a higher level of detail.

**Fig. 9.**
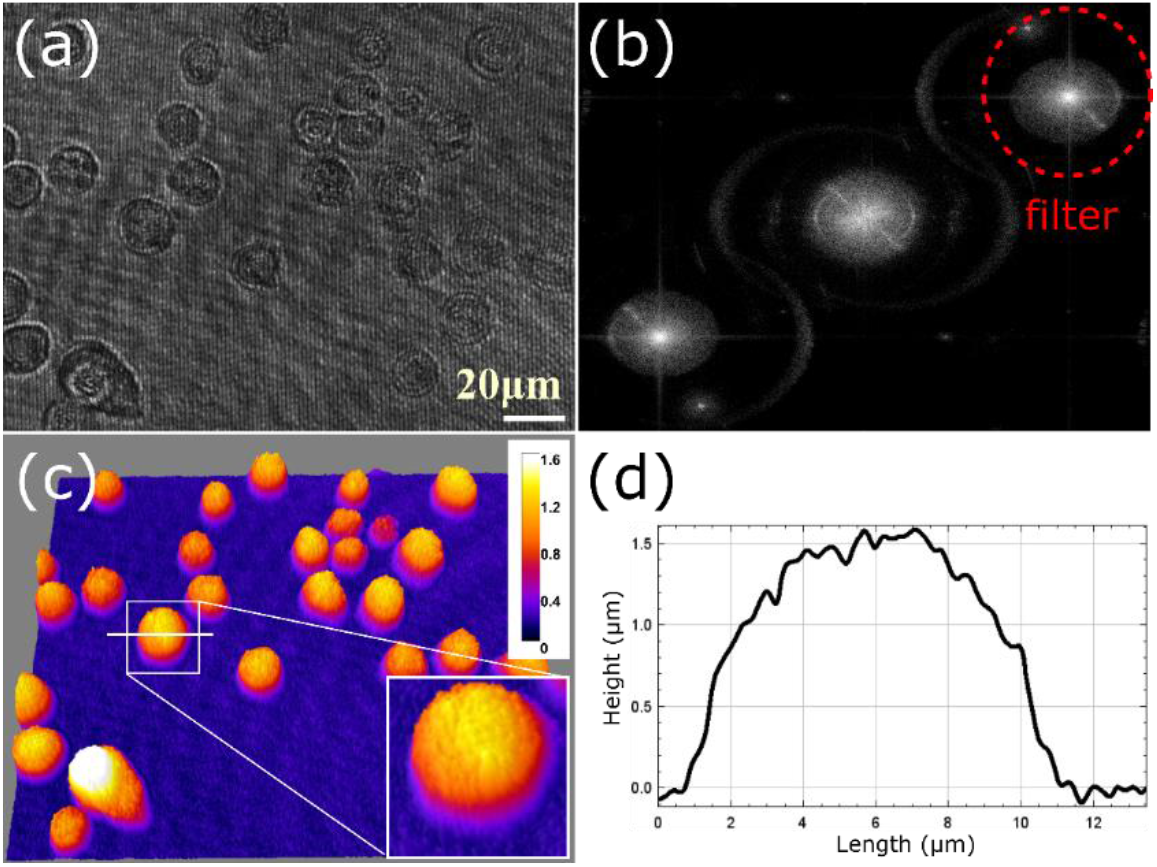
Optimized TDHM result of HeLa cells. According to the proposed methods, the optimized +1 order position ensured the largest FFT filter size ((a) and (b)). The height map (c) and cross-section plot (d) include a high level of detail. The refractive index of the HeLa cell is assumed to be 1.33, close to PBS [32]. The unit of the color scale bar in (c) is and the unit in (d) is μm. The size of the zoomed-in Region of Interest is 13 × 13*µm*.

Our comprehensive alignment procedures can be summarized as follows: the camera sensor attributes are first obtained towards the calculation of the optimal +1 order position via camera resolution; theoretical values of the object-reference beam angle and camera rotation angle are calculated; practical alignment work then constitutes rotating the *beam splitter 2* (**Fig. 1**) and camera one-dimensionally based on the calculation results to achieve the optimal +1 order position. The proposed method could be applied to other off-axis TDHM configurations, such as multiwavelength and multi-angle TDHM, to simplify the alignment process since the relationship between the beam angle and the corresponding Fourier spectrum remains the same for these systems.

## Acknowledgment

We thank Huawei Canada for providing financial support for this project under their Research Program Agreement program with the University of Toronto.

Thanks to members of Professor Aaron Wheeler’s group at the University of Toronto for providing the HeLa cells used in this study.

## Data availability

Data underlying the results presented in this paper is not publicly available at this time but may be obtained from the authors upon reasonable request.

## Reference

1. P. Memmolo, L. Miccio, M. Paturzo, G. D. Caprio, G. Coppola, P. A. Netti, and P. Ferraro, “Recent advances in holographic 3D particle tracking,” Adv. Opt. Photon., AOP 7, 713–755 (2015).

2. G. Coppola, P. Ferraro, M. Iodice, S. D. Nicola, A. Finizio, and S. Grilli, “A digital holographic microscope for complete characterization of microelectromechanical systems,” Meas. Sci. Technol. 15, 529–539 (2004).

3. B. Kemper, D. D. Carl, J. Schnekenburger, I. Bredebusch, M. Schäfer, W. Domschke, and G. von Bally, “Investigation of living pancreas tumor cells by digital holographic microscopy,” JBO 11, 034005 (2006).

4. V. K. Lam, T. C. Nguyen, B. M. Chung, G. Nehmetallah, and C. B. Raub, “Quantitative assessment of cancer cell morphology and motility using telecentric digital holographic microscopy and machine learning,” Cytometry Part A 93, 334–345 (2018).

5. S. Huang, W. Wang, J. Zeng, C. Yan, Y. Lin, and T. Wang, “Measurement of the refractive index of solutions based on digital holographic microscopy,” J. Opt. 20, 015704 (2017).

6. G. Nehmetallah and T. Nguyen, “Optical and Digital Aberration Compensation in DHM,” in Imaging and Applied Optics 2016 (2016), Paper JW4A.5 (Optica Publishing Group, 2016), p. JW4A.5.

7. Y. Liu, Z. Wang, J. Li, J. Gao, and J. Huang, “Total aberrations compensation for misalignment of telecentric arrangement in digital holographic microscopy,” OE 53, 112307 (2014).

8. J. Sun, Q. Chen, Y. Zhang, and C. Zuo, “Optimal principal component analysis-based numerical phase aberration compensation method for digital holography,” Opt. Lett., OL 41, 1293–1296 (2016).

9. A. I. Doblas, E. Sánchez-Ortiga, M. Martínez-Corral, G. Saavedra, and J. Garcia-Sucerquia, “Accurate single-shot quantitative phase imaging of biological specimens with telecentric digital holographic microscopy,” JBO 19, 046022 (2014).

10. E. Sánchez-Ortiga, P. Ferraro, M. Martínez-Corral, G. Saavedra, and A. Doblas, “Digital holographic microscopy with pure-optical spherical phase compensation,” J. Opt. Soc. Am. A, JOSAA 28, 1410–1417 (2011).

11. A. Doblas, E. Sánchez-Ortiga, M. Martínez-Corral, G. Saavedra, P. Andrés, and J. Garcia-Sucerquia, “Shift-variant digital holographic microscopy: inaccuracies in quantitative phase imaging,” Opt. Lett., OL 38, 1352–1354 (2013).

12. J. Gass, A. Dakoff, and M. K. Kim, “Phase imaging without 2π ambiguity by multiwavelength digital holography,” Opt. Lett., OL 28, 1141–1143 (2003).

13. R. Guo and F. Wang, “Compact and stable real-time dual-wavelength digital holographic microscopy with a long-working distance objective,” Opt. Express, OE 25, 24512–24520 (2017).

14. T. Bothe, J. Burke, and H. Helmers, “Spatial phase shifting in electronic speckle pattern interferometry: minimization of phase reconstruction errors,” Appl. Opt., AO 36, 5310–5316 (1997).

15. C. Trujillo, R. Castañeda, P. Piedrahita-Quintero, and J. Garcia-Sucerquia, “Automatic full compensation of quantitative phase imaging in off-axis digital holographic microscopy,” Appl. Opt., AO 55, 10299–10306 (2016).

16. B. Kemper, J. Kandulla, D. Dirksen, and G. von Bally, “Optimization of spatial phase shifting in endoscopic electronic speckle pattern interferometry,” Optics Communications 217, 151–160 (2003).

17. Z. Zhong, H. Zhao, L. Cao, M. Shan, B. Liu, W. Lu, and H. Xie, “Automatic cross filtering for off-axis digital holographic microscopy,” Results in Physics 16, 102910 (2020).

18. J. A. Picazo-Bueno, M. Trusiak, and V. Micó, “Single-shot slightly off-axis digital holographic microscopy with add-on module based on beamsplitter cube,” Opt. Express, OE 27, 5655–5669 (2019).

19. B. Kemper, D. Carl, A. Höink, G. von Bally, I. Bredebusch, and J. Schnekenburger, “Modular digital holographic microscopy system for marker free quantitative phase contrast imaging of living cells,” in Biophotonics and New Therapy Frontiers (SPIE, 2006), Vol. 6191, pp. 204–211.

20. J. Kühn, T. Colomb, F. Montfort, F. Charrière, Y. Emery, E. Cuche, P. Marquet, and C. Depeursinge, “Real-time dual-wavelength digital holographic microscopy for MEMS characterization,” in Optomechatronic Sensors and Instrumentation III (SPIE, 2007), Vol. 6716, pp. 64–73.

21. J. W. Goodman, “introduction to Fourier Optics,” (1968).

22. T. Kreis, Handbook of Holographic Interferometry: Optical and Digital Methods (John Wiley & Sons, 2006).

23. U. Schnars, C. Falldorf, J. Watson, and W. Jüptner, “Digital Holography,” in Digital Holography and Wavefront Sensing: Principles, Techniques and Applications, U. Schnars, C. Falldorf, J. Watson, and W. Jüptner, eds. (Springer, 2015), pp. 39–68.

24. T.-C. Poon and J.-P. Liu, Introduction to Modern Digital Holography: With Matlab (Cambridge University Press, 2014).

25. E. Sánchez-Ortiga, A. Doblas, G. Saavedra, M. Martínez-Corral, and J. Garcia-Sucerquia, “Off-axis digital holographic microscopy: practical design parameters for operating at diffraction limit,” Appl. Opt., AO 53, 2058–2066 (2014).

26. S. D. Nicola, A. Finizio, G. Pierattini, P. Ferraro, and D. Alfieri, “Angular spectrum method with correction of anamorphism for numerical reconstruction of digital holograms on tilted planes,” Opt. Express, OE 13, 9935–9940 (2005).

27. C. Buitrago-Duque and J. Garcia-Sucerquia, “Non-approximated Rayleigh–Sommerfeld diffraction integral: advantages and disadvantages in the propagation of complex wave fields,” Appl. Opt., AO 58, G11–G18 (2019).

28. C. J. R. Sheppard and M. Hrynevych, “Diffraction by a circular aperture: a generalization of Fresnel diffraction theory,” J. Opt. Soc. Am. A, JOSAA 9, 274–281 (1992).

29. T. Shimobaba, K. Matsushima, T. Kakue, N. Masuda, and T. Ito, “Scaled angular spectrum method,” Opt. Lett., OL 37, 4128–4130 (2012).

30. J. Kühn, T. Colomb, F. Montfort, F. Charrière, Y. Emery, E. Cuche, P. Marquet, and C. Depeursinge, “Real-time dual-wavelength digital holographic microscopy with a single hologram acquisition,” Opt. Express, OE 15, 7231–7242 (2007).

31. D. Carl, B. Kemper, G. Wernicke, and G. von Bally, “Parameter-optimized digital holographic microscope for high-resolution living-cell analysis,” Appl. Opt., AO 43, 6536–6544 (2004).

32. R. Janeiro, R. Flores, R. Flores, and J. Viegas, “Refractive index of phosphate-buffered saline in the telecom infrared C + L bands,” OSA Continuum, OSAC 4, 3039–3051 (2021).

33. J. Schindelin, I. Arganda-Carreras, E. Frise, V. Kaynig, M. Longair, T. Pietzsch, S. Preibisch, C. Rueden, S. Saalfeld, B. Schmid, J.-Y. Tinevez, D. J. White, V. Hartenstein, K. Eliceiri, P. Tomancak, and A. Cardona, “Fiji: an open-source platform for biological-image analysis,” Nat Methods 9, 676–682 (2012).

34. A. Faridian, D. Hopp, G. Pedrini, W. Osten, U. Eigenthaler, and M. Hirscher, “Different approaches to increase resolution in DHM,” in 2010 9th Euro-American Workshop on Information Optics (2010), pp. 1–3.

35. P. Gao and C. Yuan, “Resolution enhancement of digital holographic microscopy via synthetic aperture: a review,” gxjzz 3, 105–120 (2022).

36. M. K. Kim, “Principles and techniques of digital holographic microscopy,” SR 1, 018005 (2010).

